# Rich polymorphic variants of alpha satellite 34mer higher order repeats in hg38 assembly of human chromosome Y

**DOI:** 10.1101/768861

**Authors:** Ines Vlahović, Matko Glunčić, Vladimir Paar

## Abstract

A challenging problem in human population genetics is related to the unique role of human Y chromosome, with properties that distinguish humans from other species. Centromeres in primate genomes are constituted of tandem repeats of ∼ 171 bp alpha satellite monomers, commonly organized into higher order repats (HORs). Because of gaps in DNA sequencing, HOR regions as genomic “black holes” have been understudied in spite of crucial importance. Only recently the sequencing of more complete satellite DNAs becomes accessible. In human Y chromosome the largest alpha satellite higher order repeat unit 34/36mer was found, but its polymorphic variants were not investigated. Here, we study the human Y chromosome centromeric genomic sequence from hg38 assembly using our novel ALPHAsub algorithm for simple identification of alpha satellite arrays and robust GRM algorithm for HOR identification in repeat sequences. We determine the monomer alignment scheme for alpha satellite HOR array based on canonical 34mer HOR, discovering a wealth of novel polymorphic variants which include the HOR-type monomer duplications, monomer deletions/insertions or rearrangements and non-HOR insertions.

**Author Summary:** The centromere is important for segregation of chromosomes during cell division in eukaryotes. Its destabilization results in chromosomal missegregation, aneuploidy, hallmarks of cancers and birth defects. In primate genomes centromeres contain tandem repeats of ∼ 171 bp alpha satellite DNA, commonly organized into higher order repeats (HORs). In this work, we used our bioinformatics algorithms to study the human Y chromosome centromeric genomic sequence and we discover a wealth of novel polymorphic variants which include the HOR-type monomer duplications, monomer deletions/insertions or rearrangements and non-HOR insertions. These results could help to understand the role of alpha satellites and alpha HOR structures in centromeric organization and function, in particular their role in creating a functional kinetochore that is crucial for chromosome segregation during cell division.

## Introduction

It was noted that “the properties of the Y chromosome read like a list of violations of the rulebook of human genetics” and “it seems the more we know about the Y chromosome, the more questions we have” [1]. Studies of atypical structure of human Y chromosome were largely focused on gene related content [2]. On the other hand, human Y chromosome is replete with pronounced noncoding repetitive sequences [3-7].

Centromeres of all human chromosomes consist of tandem repeats of alpha satellite monomers, commonly organized as higher order repeats (HORs) superimposed on approximately periodic tandem of alpha satellite monomers [8-12]. Because of gaps in DNA sequencing, these HOR regions, like genomic “black holes” [13, 14], have been understudied in spite of their crucial importance [15, 16]. With an impressive recent progress in sequencing technology [4, 16-18], the study of more complete satellite DNAs becomes accessible [4, 19].

An alpha satellite HOR in human chromosome Y was found previously by restriction map estimates [3]. It is organized into tandemly repeating units, most of which are approximately 5.7 kb long while some variant units are about 6.0 kb long. Both the 5.7 kb and the 6.0 kb HOR units were found to consist of tandemly repeating alpha satellite monomers, with the 6.0 kb unit containing two more monomers compared to 5.7 kb unit. Later, a value of 5.941 kb was reported for this HOR unit length [2].

Sequence contigs spanning junctions at the edges of the centromere array, becoming available, enabled more extensive bioinformatics analyses of repeat patterns in human genome [13, 20-24]. However, major gaps remained in the centromeric regions of human chromosomes, as “black holes” in genomes [13, 14, 25, 26].

Using GRM algorithm, the alpha satellite HORs were identified and analysed bioinformatically for previous incomplete and gapped Build 37.1 assembly for human and Build 2.1 for chimpanzee Y chromosome [27]. The human genome reference sequence was incomplete owing to the challenge of assembling long tracts of near-identical tandem repeats in centromeres.

An alpha satellite reference model has been recently produced and incorporated in the hg38 human genome assembly [28-30]. In the hg38 human genome assembly, centromere gaps have been filled by alpha satellite reference models, which are statistical representations of homogeneous HOR arrays [31]. Only recently, a long-read strategy was applied to a human centromere. A nanopore sequencing strategy was used to generate high-quality reads that span highly repetitive DNA in centromere of one individual human Y chromosome [4]. The DYZ3 array was assembled and characterized as HOR with 5.8 kb consensus sequence without repeat inversions [4]. Instances of 6.0 kb HOR structural variant were detected, evidence for seven 6.0 kb copies within DYZ3 array was found, present in two clusters separated by 110 kb, in accordance with predictions by previous restriction map estimates [32].

## Results

Using robust ALPHAsub+GRM algorithm [27, 33, 34] we identify and analyse alpha satellite HOR arrays in hg38 assembly of Y chromosome (RefSeq Accession NC_000024.10). The resulting alpha satellite HOR ideogram for centromeric region of hg38 assembly of Y chromosome is shown, with three HOR domains I-III (Fig. 1).

**Fig. 1.**
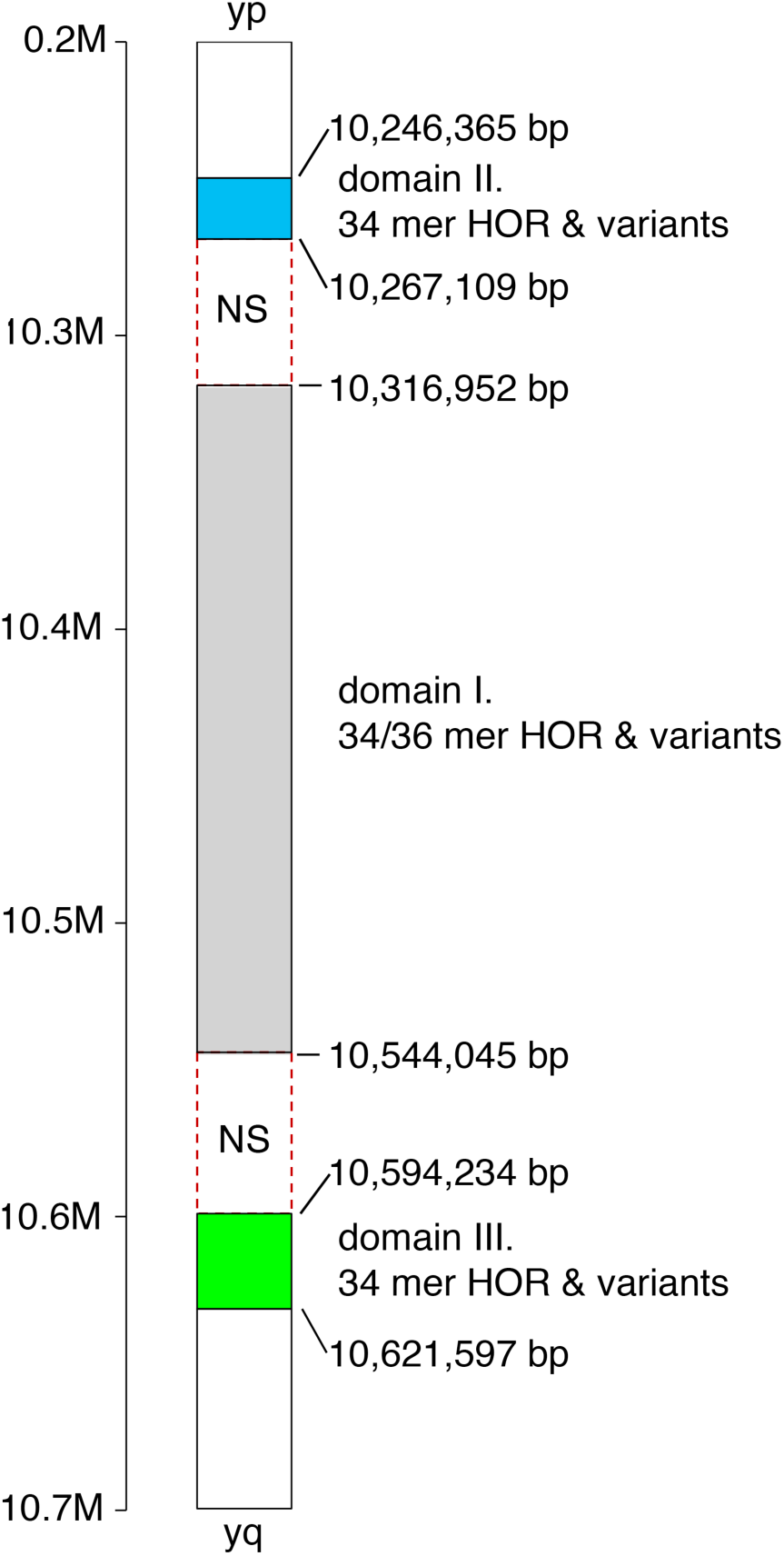
Ideogram of alpha satellite HOR arrays in domains I.-III. of hg38 assembly in centromeric/pericentromeric region of human chromosome Y. Enumeration of HOR array positions refers to hg38 assembly. NS denotes gaps in hg38 assembly.

The alpha satellite arrays extracted in the first step are given in Table 1. The GRM algorithm was extended by introducing the method ALPHAsub used to identify positions of alpha satellite arrays in DNA sequence, regardless weather they are of HOR or nonHOR type. In this way we determined segments with alpha satellite arrays. We have found 28 different alpha satellite arrays in human chromosome Y, three of them arranged in 34mer/36mer HOR structures. Previously, for human chromosome Y, only the 5.7 kb and the 6.0 kb alpha satellite HOR units were found[2-4].

**Table 1.**
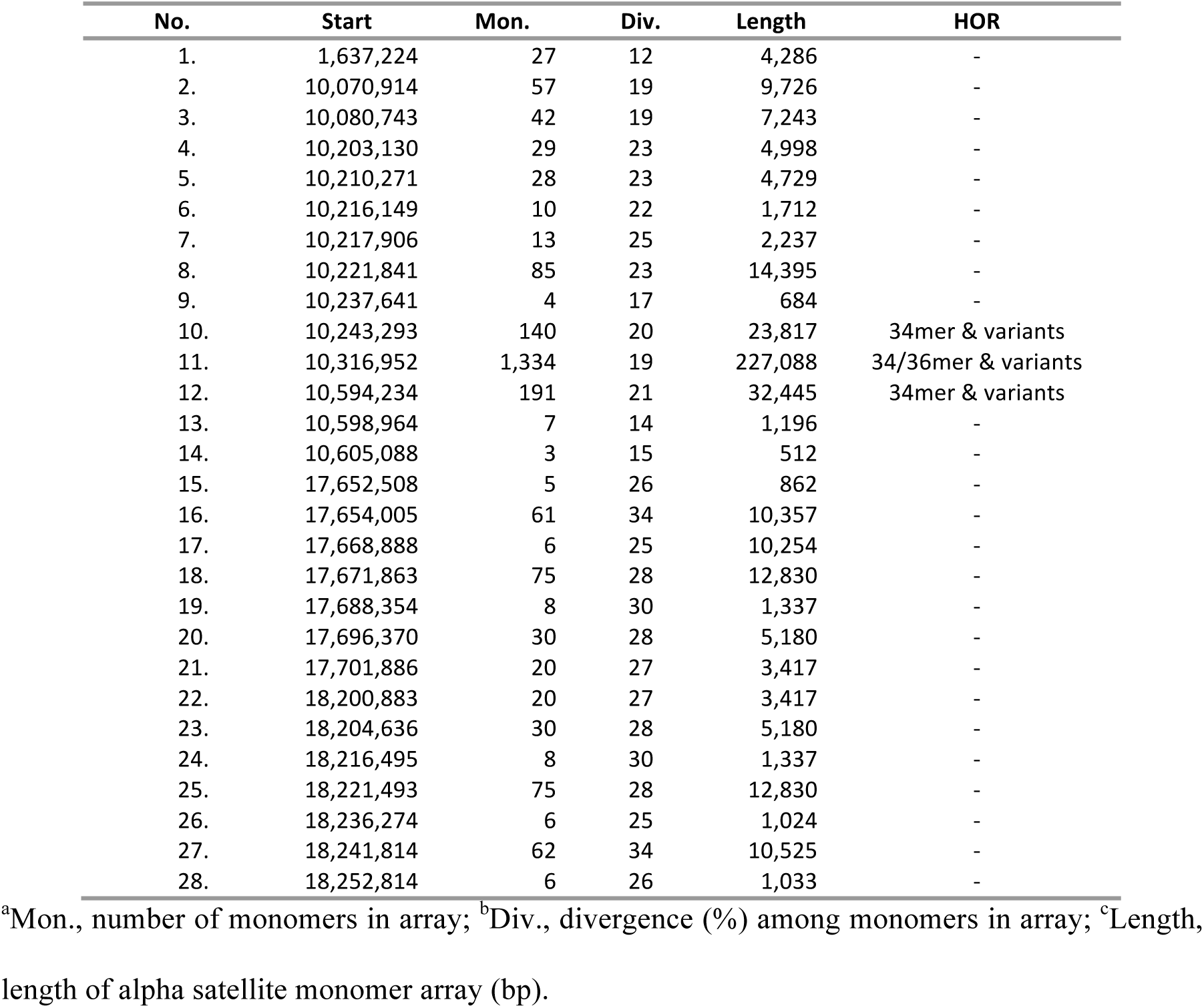
Alpha satellite arrays in centromeric/pericentromeric region of hg38 sequence of human chromosome Y obtained using ALPHAsub algorithm.

| No. | Start | Mon. | Div. | Length | HOR |
| --- | --- | --- | --- | --- | --- |
| 1. | 1,637,224 | 27 | 12 | 4,286 | - |
| 2. | 10,070,914 | 57 | 19 | 9,726 | - |
| 3. | 10,080,743 | 42 | 19 | 7,243 | - |
| 4. | 10,203,130 | 29 | 23 | 4,998 | - |
| 5. | 10,210,271 | 28 | 23 | 4,729 | - |
| 6. | 10,216,149 | 10 | 22 | 1,712 | - |
| 7. | 10,217,906 | 13 | 25 | 2,237 | - |
| 8. | 10,221,841 | 85 | 23 | 14,395 | - |
| 9. | 10,237,641 | 4 | 17 | 684 | - |
| 10. | 10,243,293 | 140 | 20 | 23,817 | 34mer & variants |
| 11. | 10,316,952 | 1,334 | 19 | 227,088 | 34/36mer & variants |
| 12. | 10,594,234 | 191 | 21 | 32,445 | 34mer & variants |
| 13. | 10,598,964 | 7 | 14 | 1,196 | - |
| 14. | 10,605,088 | 3 | 15 | 512 | - |
| 15. | 17,652,508 | 5 | 26 | 862 | - |
| 16. | 17,654,005 | 61 | 34 | 10,357 | - |
| 17. | 17,668,888 | 6 | 25 | 10,254 | - |
| 18. | 17,671,863 | 75 | 28 | 12,830 | - |
| 19. | 17,688,354 | 8 | 30 | 1,337 | - |
| 20. | 17,696,370 | 30 | 28 | 5,180 | - |
| 21. | 17,701,886 | 20 | 27 | 3,417 | - |
| 22. | 18,200,883 | 20 | 27 | 3,417 | - |
| 23. | 18,204,636 | 30 | 28 | 5,180 | - |
| 24. | 18,216,495 | 8 | 30 | 1,337 | - |
| 25. | 18,221,493 | 75 | 28 | 12,830 | - |
| 26. | 18,236,274 | 6 | 25 | 1,024 | - |
| 27. | 18,241,814 | 62 | 34 | 10,525 | - |
| 28. | 18,252,814 | 6 | 26 | 1,033 | - |

<sup>a</sup>Mon., number of monomers in array; <sup>b</sup>Div., divergence (%) among monomers in array; <sup>c</sup>Length, length of alpha satellite monomer array (bp).

The GRM peak at 5786 bp corresponding to the major 34mer/36mer HOR in GRM diagram is sizable (Fig. 2), because of large number of approximately regular HOR copies in HOR array. Applying ALPHAsub+GRM algorithm we determined the monomer alignment schemes for alpha satellite HOR arrays in hg38 assembly for human Y chromosome in domains I, II, and III (Fig. 3a-c, respectively). Going along genome sequence, the monomers are positioned in the monomer alignment scheme in order of appearance. In constructing monomer alignment scheme monomers are assigned to the same monomer type if divergence is less than 5 % and are placed in the same column of the scheme. Otherwise, monomers do not belong to types included in HOR and are referred to as non-HOR monomers.

**Fig. 2.**
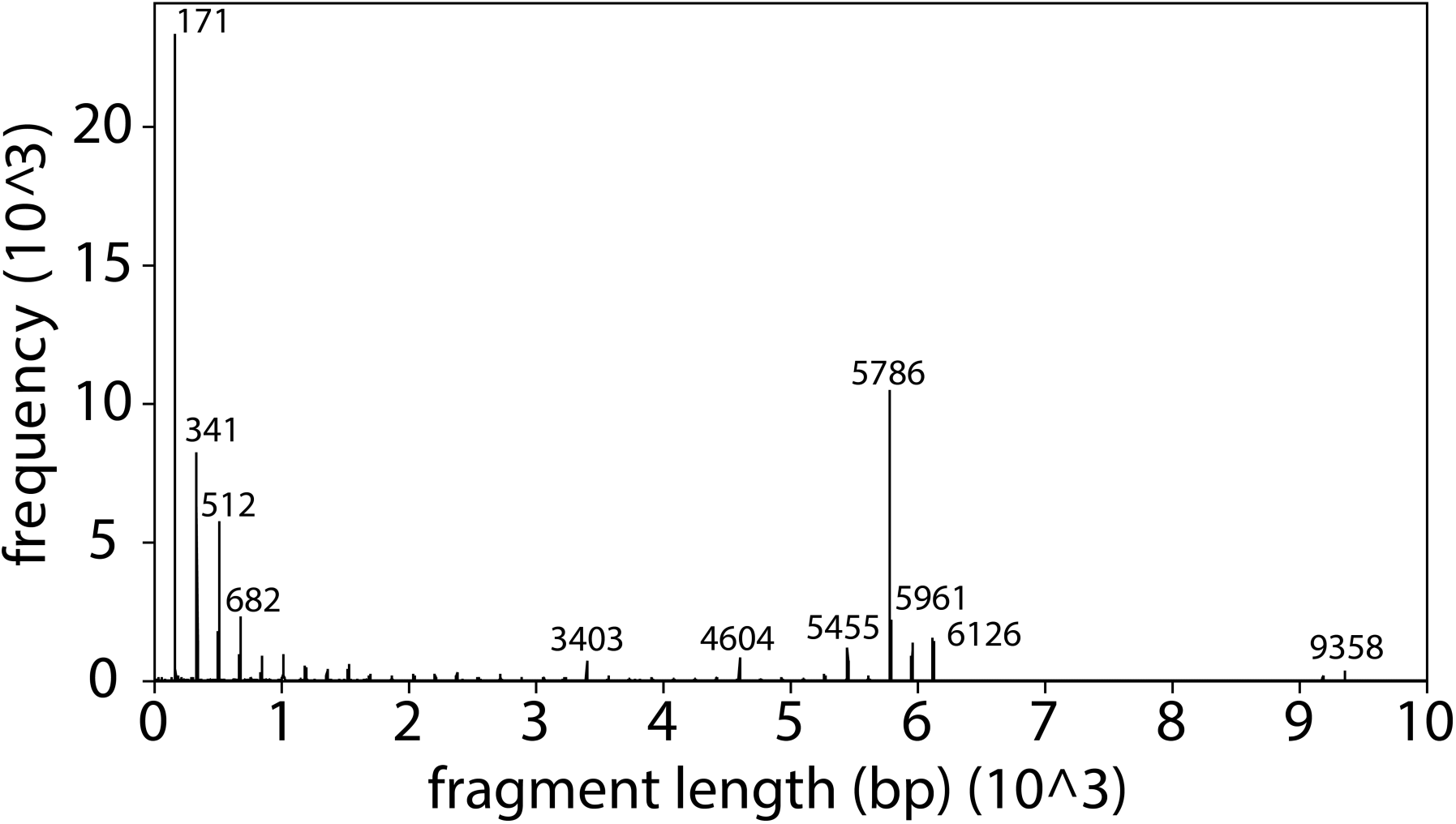
GRM diagram for domain I: of hg38 assembly of human chromosome Y. The pronounced peak at 5,786 bp corresponds to 34/36mer HOR. Since the average length of alpha satellite monomer is ∼ 171 bp, the 5,786 bp peak in GRM diagram of human Y chromosome corresponds to *n* ∼ 5,786 bp / 171 bp ∼ 33.8 ∼ 34 monomers. This is close to the previous length estimates of 5.7 kb [3] and 5.8 kb [4] for the major HOR in human chromosome Y.

**Fig. 3.**
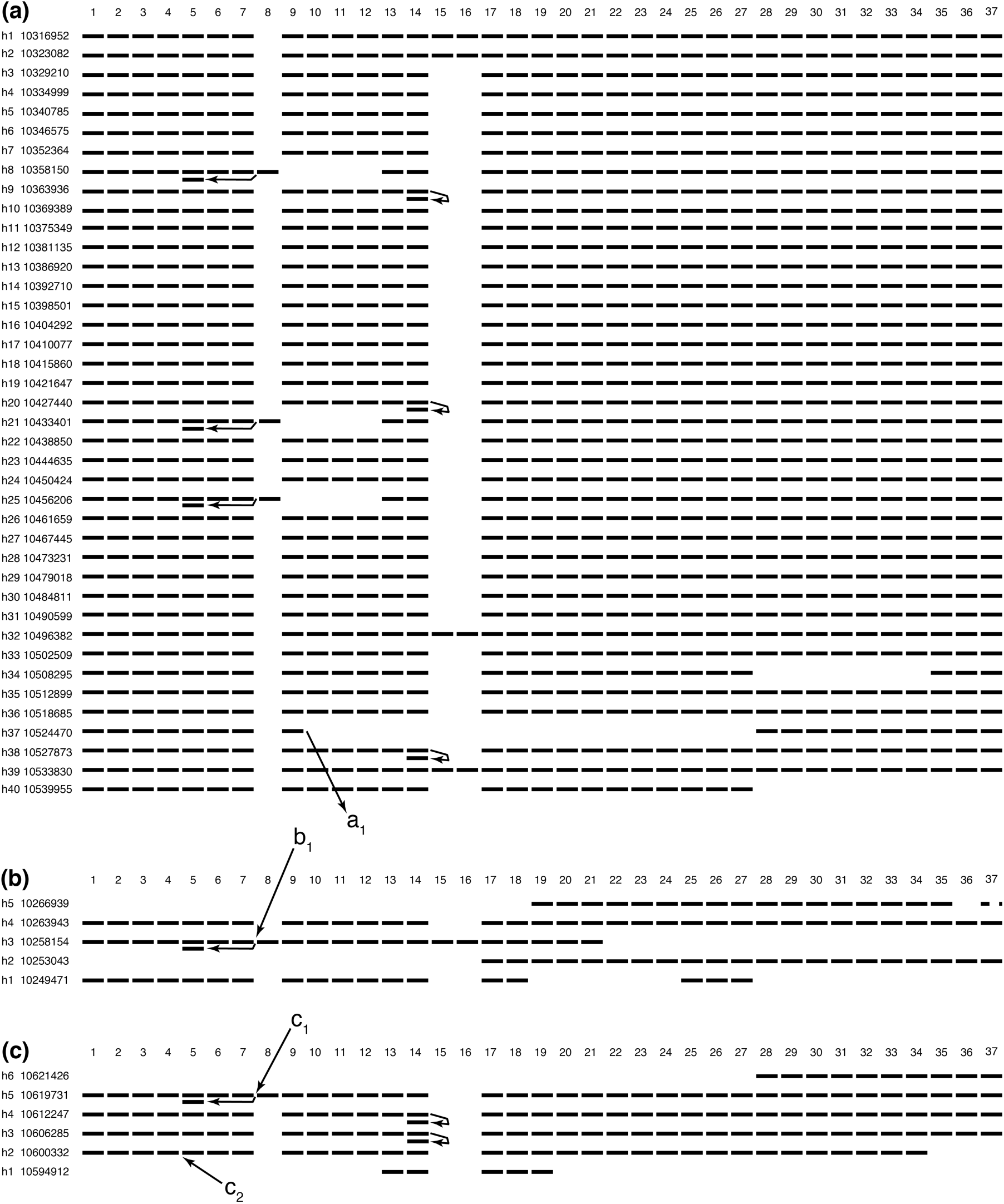
Monomer alignment schemes for canonical alpha satellite 34mer HORs and its polymorphic variants in hg38 assembly of human Y chromosome – (a) domain I, (b) domain II, and (c) domain III. Top row in all three schemes denotes types of alpha satellite monomers. Each monomer is schematically presented by a horizontal bar in the column of the corresponding monomer type. Any duplicate of a HOR monomer (in most cases defined by divergence of less than 5%) is displayed in alignment scheme by an additional horizontal bar below the bar of the corresponding primary monomer; a position of a duplicate monomer is indicated by an arrow. 37 monomers m1, m2, … m37 denote the 37 types of monomers in order of appearance. The monomer types m8, m15 and m16 are absent in most of HOR copies and therefore, they are referred to as non-canonical monomers. Thus, most of HOR copies, like for example h5 (10340785), contain 34 monomers m1, m2, …, m7, m9, m10…, m14, m17, m18, … m37, representing the canonical 34mer HOR. The domains II and III have been inserted in the hg38 assembly sequence of Y chromosome in reverse orientation, considering the orientation of domain I. Here, we present domain II and domain III monomer alignment schemes for canonical alpha satellite 34mer HORs and its polymorphic variants in direct orientations and adjust the start of HOR sequences to match the individual monomers from domains II and III to monomers in domain I. In addition, each HOR structure contains additional inserted non-HOR alpha satellite monomers: a_1_ - array of two inserted monomers, b_1_ - array of eight inserted monomers, c_1_ - array of eight inserted monomers, and c_2_ - one inserted monomer. The arrays of eight inserted monomers b_1_ and c_1_ are similar up to 5% (see Fig. 4d).

The mean divergence among monomers within each HOR copy in domain I is ∼ 20%, and the mean divergence among monomers of the same type in different HOR copies is ∼ 0.5%. For 34mer HOR array in domain II the mean divergence is ∼ 20 % and ∼ 3 %, respectively; and for 34mer HOR array in domain III ∼ 20 % and ∼ 1 %, respectively, not counting monomer insertions/deletions. The corresponding consensus sequences of monomers in 34/36mer are given in Supplementary Table 1.

### 34/36mer HOR and its polymorphic variants in domain I

Using ALPHAsub+GRM algorithm in domain I of hg38 assembly, the 40 HOR copies are obtained: 27 complete 34mer HOR copies (referred to as canonical) and 13 polymorphic variants (Fig. 3a). The monomer types m8, m15 and m16 are absent in most of HOR copies and thus they are referred to as non-canonical monomers. In the aligned monomer scheme each horizontal bar presenting a monomer is characterized by its monomer type m1, m2, m37 and arrays of non-HOR monomers are characterized by arrow and symbol of insertion (for example a_1_ in Fig. 3a). For example, the HOR copy h1 consists of 36 monomer tandems of types m1-m7 and m9-m37 (displayed by horizontal bars No. 1-7 and 9-37). HOR copy h37 consists of a 20 monomers, tandem m1-m7 (monomers No. 1-7), monomer m9 (No. 9), tandem m28-m37 (monomers No. 28-37) and two non-HOR monomer array a_1_ inserted after monomer m9.

Four polymorphic variants are complete 36mers, three 32mers, three 35mers, one 27mer, one 24mer, and one 20mer (Table 2a). The four complete variant 36mer HOR copies arise from canonical 34mer by inserting after m14 two additional monomers. Such are HOR copies h1, h2, h32, h39. This HOR array, containing both 34mer and 36mer copies, is referred to as 34/36mer HOR. Occasional monomer duplications, similar as found here, appear also in alpha satellite HORs in some other human chromosomes (for example, [27, 35]).

**Table 2.** Canonical alpha satellite 34mer HOR and its polymorphic variants in domains I-III of hg38 assembly of human chromosome Y.

| *n*mer | HOR<br>copies | monomer<br>del. | ins. | HOR copy |
| --- | --- | --- | --- | --- |
| (a) domain I |  |  |  |  |
| 34mer | 27 |  |  | h3-h7, h10-h19, h22-h24,<br>h26-h31, h33, h35-h36 |
| 36mer | 4 |  | 2 | h1-h2, h32, h39 |
| 35mer | 3 |  | 1 | h9, h20, h38 |
| 32mer | 3 | 4 | 2 | h8, h21, h25 |
| 27mer | 1 | 7 |  | h34 |
| 24mer | 1 | 10 |  | h40 |
| 20mer | 1 | 16 | 2 | h37 |
| (b) domain II |  |  |  |  |
| 34mer | 1 |  |  | h4 |
| 28mer | 1 | 16 | 12 | h3 |
| 21mer | 1 | 13 |  | h2 |
| 18mer | 2 | 16 |  | h1, h5 |
| (c) domain III |  |  |  |  |
| 35mer | 2 |  | 1 | h3, h4 |
| 44mer | 1 |  | 10 | h5 |
| 32mer | 1 | 3 | 1 | h2 |
| 10mer | 1 | 24 |  | h6 |
| 5mer | 1 | 29 |  | h1 |

del = repeat monomer deletions expressed with respect to canonical 34mer; ins = repeat and non-repeat monomer insertions.
Number *n* of monomers in *n*mer HOR implies the number of different types of monomers present in HOR copy.

The variant 35mer HOR copies h9, h20 and h38 arise from canonical 34mer HOR copy by duplicating the monomer m14, resulting in the variant monomer sequence … m13 m14 m14 m17 m18 … This variant triplet of HOR copies characterizes domain I and domain III. It is noted that monomer duplications appear in alpha satelite HORs also in other human chromosomes (for example, [27, 35]). Monomer duplication is present in six variant HOR copies for domain I. A particular case is combined monomer duplication, deletion and insertion in HOR copies h8, h21 and h25. In these three variant 32mer HOR copies the typical monomers m1-m7, m13-m14 and m17-m37 are intact. Then m5 is duplicated, a non-canonical monomer m8 is inserted and typical m9-m12 monomers are absent. These three HOR copies appear as polymorphic variants, where the 5-monomer region m8-m12 is distorted in a specific way, including replacement by one duplicate and one atypical monomer. This results in variant monomer sequence with … m4, m5, m6, m7, m5, m8, m13, m14, m17…

Variant HOR copies h34 and h37 involve significant deletions of monomers, and h40 is located at the end of domain I and is missing last ten monomers (m28-m37). Variant HOR copy h37 also involve insertion of two non-HOR alpha satellite monomers (non-repeating monomers which don’t have another copies within HOR structure)

### 34mer HOR and its polymorphic variants in domain II

The domains II and III have been inserted in the hg38 assembly sequence of Y chromosome in reverse orientation, considering the orientation of domain I. We have presented domain II and domain III monomer alignment schemes for canonical alpha satellite 34mer HORs and its polymorphic variants in direct orientations (Fig 3b and 3c) and we have conveniently adjust the start of HOR sequences to match the individual monomers from domains II and III to monomers in domain I (Fig 4a and 4b).

**Fig. 4.**
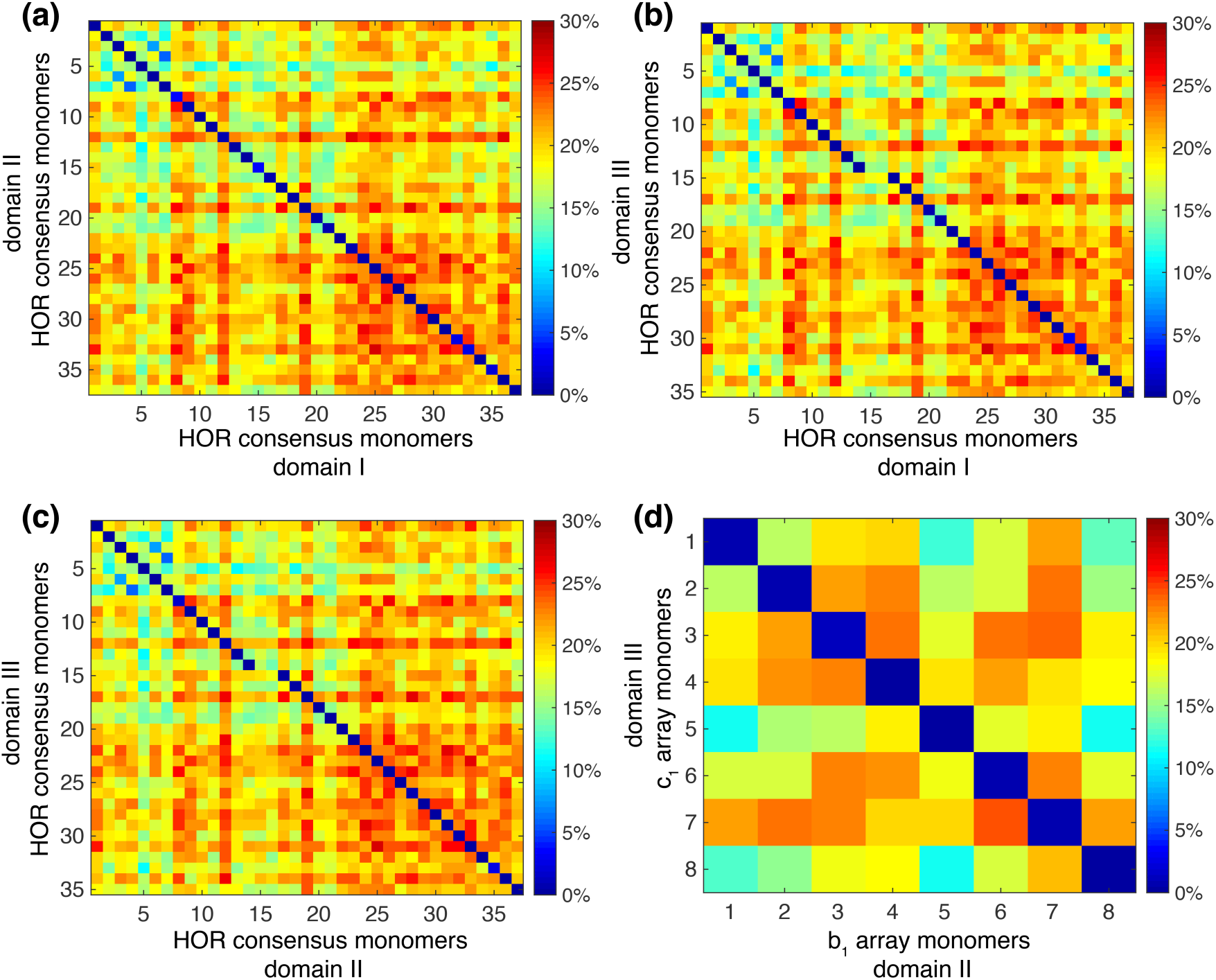
Heatmap for divergence of consensus monomer types among domains: (a) HOR consensus monomers in domains I vs. II, (b) HOR consensus monomers in domains I vs. III, (c) HOR consensus monomers in domains II vs. III, (d) non-HOR monomer array in domain II vs III (b1 vs c1 from Fig 3). We conveniently adjust the starting monomers within consensus HOR copies. The positions of the same type monomers in domains I vs. II, I vs. III, and II vs. III lie on diagonal. It is obvious that monomers m15 and m16 in domains I and II have no monomer counterparts (below 5% identity) in domain III which corresponds the HOR schemes from Fig 3. The array of 8 inserted monomers in domain II (b_1_ in Fig 3b) is similar (up to 5%) to array of 8 inserted monomers in domain III (c_1_ in Fig 3c).

In domain II we identified segments classified in five 34mer HOR copies h1-h5 (Fig. 3b and Table 2b). They contain 45 different types of alpha satellite monomers: out of 34 canonical and 3 non-canonical monomers (m8, m15, m16) from domain I, all have counterparts in domain II. The corresponding similarity of monomer types between domains I vs. II are shown in Fig 4a. The other monomers are insertions of non-HOR monomers.

In this HOR array each of 34 HOR-monomers m1-m34 from domain I is repeated, most of them threefold. However, besides these HOR-repeating monomers, the HOR copy h3 has some inserted non-HOR monomers, appearing only once in the aligned scheme, labelled as b_1_ (array of 8 non-HOR monomers). This array of 8 inserted monomers is similar (up to 5%) to array of 8 inserted monomers in domain III (labelled as c_1_) (Fig. 4d). This array are followed by a duplication of monomer m5, then by an insertion of non-canonical monomer m8, then by a tandem of HOR monomers m9-m4, then by an insertion of two non-canonical monomers m15 and m16, and then by a tandem of HOR monomers m17-m37.

### 34mer HOR and its polymorphic variants in domain III

Using ALPHAsub+GRM algorithm the scheme of aligned monomer structure for HORs in domain III with 6 HOR copies was determined (Fig. 3c and Table 2c). The corresponding consensus HOR unit is almost identical to consensus HOR unit in domain I (divergence ∼ 0.3 %,). The corresponding similarity of monomer types between domains I vs. III are shown in Fig 4b.

All six HOR copies in domain III are polymorphic variants of canonical 34mer HOR. Two HOR copies (h3 and h4) are 35mers, where both copies have one monomer duplicated (m14, like 35mers in domain I). The HOR copy h2 has one non-HOR monomer insertion (marked as insertion c_2_ in Fig. 3c) and three monomers deletion (m35-m37). The HOR copy h5 contains 44 HOR monomer types, and is sizably distorted by addition of array of 8 non-HOR monomers (labelled as c_1_ in Fig 3c) that follow after m7 and are continued by non-canonical monomer insertion m8. These additional 8 monomers diverge from all classical HOR monomers by more than 5% and are similar to 8 additional monomers from domain II (Fig 4d). The structure of HOR copies h1 (5mer) and h6 (10mer) are determined by their location, the start and the end of domain III, respectively.

Full monomer divergence matrices between consensus monomers from domains I vs. II, I vs. III, and II vs. III are shown by heatmap in Fig. 4. As could be predicted from Fig 3, m15 and m16 in domains I and II have no monomer counterparts (below 5% identity) in domain III (Fig 4b and 4c).

## Discussion

Recent rapidly improving second and third generation sequencing opens the possibility to determine complete ensemble of alpha satellite HORs in the whole human genome, which will enable broader investigations of alpha satellite HORs, their polymorphic variants and their influence on centromere dynamics. Previously, in chromosome Y, HOR with 5.8 kb consensus sequence and 6.0 kb HOR structural variant were detected [3, 4] that correspond to 34mer and 36mer HORs, respectively. In this paper, we have discovered a wealth of novel polymorphic variants, which include the HOR-type monomer duplications, monomer deletions/insertions or rearrangements and non-HOR insertions. In particular, these polimorfic varyants result with HOR structures up to 44 monomers length. These results could help to understand the role of alpha satellites and alpha HOR structures in centromeric organization and function, in particular their role in formation of functional kinetochore. One could expect that rich long HOR repeat units will be found also in centromere of some other human chromosomes. The coming years may bring exciting new developments in HOR investigations.

## Methods

In this study the hg38 assembly sequence of Y chromosome (RefSeq Accession.version NC_000024.10) was used for HOR analysis downloaded from: ftp://ftp.ncbi.nlm.nih.gov/genomes/H_sapiens/ARCHIVE/ANNOTATION_RELEASE.1_08/Assembled_chromosomes/seq/.

Here we used our robust computational algorithm GRM - Global Repeat Map algorithm[27, 33, 34] convenient for HOR identification in novel centromeric repeat sequences and an ALPHAsub algorithm convenient for simple identification of alpha satellite arrays.

### ALPHAsub algorithm

ALPHAsub algorithm is a simple method for extraction of alpha satellite tandem arrays from a given genomic sequence, irrespectively of whether they are organized into HORs or not. As a convenient “ideal key word” (a “seed”), we use a robust 28-bp segment from alpha satellite DNA sequences, TGAGAAACTGCTTTGTGATGTGTGCATT and its reverse complement. This choice of a well conserved region of known alpha satellite tandem is based on our previous experience with alpha satellite tandem arrays [27, 33, 35]. First, using the Levenshtein distance algorithm [36], all positions in the whole chromosome are determined where the 28-bp sequence of “ideal key word” or its reverse complement differs from a “real key word” by at most nine nucleotides. Second, the distances between positions of neighbouring “real key words” are calculated. Third, only those “real key words” are retained for which distance to its previous neighbour is approximately equal to 171 bp or to a multiple of 171 bp (*d* (*n, n* − 1)∼*m* · 171; *m =* 1, 2, …). In the latter case (*m* > 1), the additional “real key words” (one for *m =* 2, two for *m =* 2, and so on) are determined in the sequence between “real key word” and its previous neighbour, using the Levenshtein distance algorithm, at positions with the smallest difference of “real key words” compared to “ideal key word” or its reverse complement. In general, a distance between the additional “real key words”, obtained by this method, is always approximately equal to 171 bp. In this way, we determined positions of all alpha satellites within chromosome Y. In the next step, using positions of “real key words”, all alpha satellites from hg38 DNA sequence for chromosome Y are extracted and different alpha satellite ensembles are identified. On this basis, we have designed our ALPHAsub algorithm and computer program. Applying ALPHAsub program to the hg38 sequence of Y chromosome we determine location of all alpha satellite arrays within genomic sequence. In this way we determine regions within Y chromosome that contain alpha satellites arrays.

### GRM algorithm

Global repeat algorithm (GRM) is an efficient and robust novel method to identify and study repeats, especially HORs, in a given DNA sequence [27, 33, 34]. For long DNA sequences of whole chromosomes, the noise in GRM diagram increases with increasing length of HOR repeat unit. This noise is significantly reduced by applying GRM to those regions which contain alpha satellite arrays selected using ALPHAsub algorithm for analysis of the whole chromosome sequence. We note that the GRM algorithm chooses the starting point autonomously, causing a difference of starting point with respect to standardly used sequence of consensus monomer. This choice of starting point does not influence the results.

### ALPHAsub+GRM algorithm: GRM algorithm expanded by ALPHAsub algorithm

Successive application of ALPHAsub and GRM algorithms is used for identification and analysis of alpha satellite HORs in a whole chromosome sequence: in the first step we identify chromosome regions that contain alpha satellite arrays and in the second step we perform GRM computation for these regions. The algorithm is freely available on our web server genom.hazu.hr at https://genom.hazu.hr/tools.html. For identification of higher order structures, we use Needleman-Wunsch algorithm that creates a divergence matrix where diagonals highlight higher order structures of *n*-mers. We also show divergences in heatmap graphs.

## Acknowledgements

We thank C. Tyler-Smith for stimulating our interest for alpha satellites. We also acknowledge support from the QuantiXLie Centre of Excellence, a project cofinanced by the Croatian Government and European Union through the European Regional Development Fund - the Competitiveness and Cohesion Operational Programme (Grant KK.01.1.1.01.0004), and the grant IP-2014-09-3626 from Croatian Science Foundation.

## Author Contributions

I.V. and M.G. performed the computations. M.G. wrote computational algorithm ALPHAsub. V.P. supervised the study. All authors analysed computational results. V.P. wrote the manuscript. All authors read and approved the final version of the manuscript.

## Supporting information captions

### Supplementary Tables S1

**a) Consensus sequence of canonical 34/36mer HOR in domain I**

**b) Consensus sequence of variant 34mer HOR in domain II**

**c) Consensus sequence of variant 34mer HOR in domain III**

## References

1. Jobling MA, Tyler-Smith C. The human Y chromosome: an evolutionary marker comes of age. Nat Rev Genet. 2003;4(8):598–612. doi: 10.1038/nrg1124. PubMed PMID: 12897772.

2. Skaletsky H, Kuroda-Kawaguchi T, Minx PJ, Cordum HS, Hillier L, Brown LG, et al. The male-specific region of the human Y chromosome is a mosaic of discrete sequence classes. Nature. 2003;423(6942):825–37. doi: 10.1038/nature01722. PubMed PMID: 12815422.

3. Tyler-Smith C, Brown WR. Structure of the major block of alphoid satellite DNA on the human Y chromosome. J Mol Biol. 1987;195(3):457-70. PubMed PMID: 2821279.

4. Jain M, Olsen HE, Turner DJ, Stoddart D, Bulazel KV, Paten B, et al. Linear assembly of a human centromere on the Y chromosome. Nat Biotechnol. 2018;36(4):321–3. doi: 10.1038/nbt.4109. PubMed PMID: 29553574; PubMed Central PMCID: PMCPMC5886786.

5. Tyler-Smith C, Oakey RJ, Larin Z, Fisher RB, Crocker M, Affara NA, et al. Localization of DNA sequences required for human centromere function through an analysis of rearranged Y chromosomes. Nat Genet. 1993;5(4):368–75. doi: 10.1038/ng1293-368. PubMed PMID: 8298645.

6. Rozen S, Skaletsky H, Marszalek JD, Minx PJ, Cordum HS, Waterston RH, et al. Abundant gene conversion between arms of palindromes in human and ape Y chromosomes. Nature. 2003;423(6942):873–6. doi: 10.1038/nature01723. PubMed PMID: 12815433.

7. Perry GH, Tito RY, Verrelli BC. The evolutionary history of human and chimpanzee Y-chromosome gene loss. Mol Biol Evol. 2007;24(3):853–9. doi: 10.1093/molbev/msm002. PubMed PMID: 17218643.

8. Manuelidis L. Chromosomal localization of complex and simple repeated human DNAs. Chromosoma. 1978;66(1):23-32. PubMed PMID: 639625.

9. Willard HF. Chromosome-specific organization of human alpha satellite DNA. Am J Hum Genet. 1985;37(3):524-32. PubMed PMID: 2988334; PubMed Central PMCID: PMCPMC1684601.

10. Jorgensen AL, Bostock CJ, Bak AL. Homologous subfamilies of human alphoid repetitive DNA on different nucleolus organizing chromosomes. Proc Natl Acad Sci U S A. 1987;84(4):1075-9. PubMed PMID: 3469648; PubMed Central PMCID: PMCPMC304364.

11. Waye JS, Willard HF. Nucleotide sequence heterogeneity of alpha satellite repetitive DNA: a survey of alphoid sequences from different human chromosomes. Nucleic Acids Res. 1987;15(18):7549-69. PubMed PMID: 3658703; PubMed Central PMCID: PMCPMC306267.

12. Aldrup-Macdonald ME, Sullivan BA. The past, present, and future of human centromere genomics. Genes (Basel). 2014;5(1):33-50. PubMed PMID: 24683489; PubMed Central PMCID: PMCPMC3966626.

13. Rudd MK, Willard HF. Analysis of the centromeric regions of the human genome assembly. Trends Genet. 2004;20(11):529–33. doi: 10.1016/j.tig.2004.08.008. PubMed PMID: 15475110.

14. Henikoff S. Near the edge of a chromosome’s “black hole". Trends Genet. 2002;18(4):165-7. PubMed PMID: 11932007.

15. Treangen TJ, Salzberg SL. Repetitive DNA and next-generation sequencing: computational challenges and solutions. Nat Rev Genet. 2011;13(1):36–46. doi: 10.1038/nrg3117. PubMed PMID: 22124482; PubMed Central PMCID: PMCPMC3324860.

16. Lower SS, McGurk MP, Clark AG, Barbash DA. Satellite DNA evolution: old ideas, new approaches. Curr Opin Genet Dev. 2018;49:70–8. doi: 10.1016/j.gde.2018.03.003. PubMed PMID: 29579574; PubMed Central PMCID: PMCPMC5975084.

17. Alkan C, Ventura M, Archidiacono N, Rocchi M, Sahinalp SC, Eichler EE. Organization and evolution of primate centromeric DNA from whole-genome shotgun sequence data. PLoS Comput Biol. 2007;3(9):1807–18. doi: 10.1371/journal.pcbi.0030181. PubMed PMID: 17907796; PubMed Central PMCID: PMCPMC1994983.

18. van Dijk EL, Jaszczyszyn Y, Naquin D, Thermes C. The Third Revolution in Sequencing Technology. Trends Genet. 2018;34(9):666–81. doi: 10.1016/j.tig.2018.05.008. PubMed PMID: 29941292.

19. McNulty SM, Sullivan BA. Alpha satellite DNA biology: finding function in the recesses of the genome. Chromosome Res. 2018;26(3):115–38. doi: 10.1007/s10577-018-9582-3. PubMed PMID: 29974361; PubMed Central PMCID: PMCPMC6121732.

20. Rudd MK, Schueler MG, Willard HF. Sequence organization and functional annotation of human centromeres. Cold Spring Harb Symp Quant Biol. 2003;68:141-9. PubMed PMID: 15338612.

21. Rosandic M, Paar V, Basar I. Key-string segmentation algorithm and higher-order repeat 16mer (54 copies) in human alpha satellite DNA in chromosome 7. J Theor Biol. 2003;221(1):29-37. PubMed PMID: 12634041.

22. Nusbaum C, Mikkelsen TS, Zody MC, Asakawa S, Taudien S, Garber M, et al. DNA sequence and analysis of human chromosome 8. Nature. 2006;439(7074):331–5. doi: 10.1038/nature04406. PubMed PMID: 16421571.

23. Gelfand Y, Rodriguez A, Benson G. TRDB--the Tandem Repeats Database. Nucleic Acids Res. 2007;35(Database issue):D80–7. doi: 10.1093/nar/gkl1013. PubMed PMID: 17175540; PubMed Central PMCID: PMCPMC1781109.

24. Warburton PE, Hasson D, Guillem F, Lescale C, Jin X, Abrusan G. Analysis of the largest tandemly repeated DNA families in the human genome. BMC Genomics. 2008;9:533. doi: 10.1186/1471-2164-9-533. PubMed PMID: 18992157; PubMed Central PMCID: PMCPMC2588610.

25. Schueler MG, Higgins AW, Rudd MK, Gustashaw K, Willard HF. Genomic and genetic definition of a functional human centromere. Science. 2001;294(5540):109–15. doi: 10.1126/science.1065042. PubMed PMID: 11588252.

26. Miga KH, Newton Y, Jain M, Altemose N, Willard HF, Kent WJ. Centromere reference models for human chromosomes X and Y satellite arrays. Genome Res. 2014;24(4):697–707. doi: 10.1101/gr.159624.113. PubMed PMID: 24501022; PubMed Central PMCID: PMCPMC3975068.

27. Paar V, Gluncic M, Basar I, Rosandic M, Paar P, Cvitkovic M. Large tandem, higher order repeats and regularly dispersed repeat units contribute substantially to divergence between human and chimpanzee Y chromosomes. J Mol Evol. 2011;72(1):34–55. doi: 10.1007/s00239-010-9401-8. PubMed PMID: 21103868.

28. Rosenbloom KR, Armstrong J, Barber GP, Casper J, Clawson H, Diekhans M, et al. The UCSC Genome Browser database: 2015 update. Nucleic Acids Res. 2015;43(Database issue):D670–81. doi: 10.1093/nar/gku1177. PubMed PMID: 25428374; PubMed Central PMCID: PMCPMC4383971.

29. Shepelev VA, Uralsky LI, Alexandrov AA, Yurov YB, Rogaev EI, Alexandrov IA. Annotation of suprachromosomal families reveals uncommon types of alpha satellite organization in pericentromeric regions of hg38 human genome assembly. Genom Data. 2015;5:139–46. doi: 10.1016/j.gdata.2015.05.035. PubMed PMID: 26167452; PubMed Central PMCID: PMCPMC4496801.

30. Tyner C, Barber GP, Casper J, Clawson H, Diekhans M, Eisenhart C, et al. The UCSC Genome Browser database: 2017 update. Nucleic Acids Res. 2017;45(D1):D626–D34. doi: 10.1093/nar/gkw1134. PubMed PMID: 27899642; PubMed Central PMCID: PMCPMC5210591.

31. Uralsky LI, Shepelev VA, Alexandrov AA, Yurov YB, Rogaev EI, Alexandrov IA. Classification and monomer-by-monomer annotation dataset of suprachromosomal family 1 alpha satellite higher-order repeats in hg38 human genome assembly. Data Brief. 2019;24:103708. doi: 10.1016/j.dib.2019.103708. PubMed PMID: 30989093; PubMed Central PMCID: PMCPMC6447721.

32. Tyler-Smith C. Structure of repeated sequences in the centromeric region of the human Y chromosome. Development. 1987;101 Suppl:93-100. PubMed PMID: 3503726.

33. Gluncic M, Paar V. Direct mapping of symbolic DNA sequence into frequency domain in global repeat map algorithm. Nucleic Acids Res. 2013;41(1):e17. doi: 10.1093/nar/gks721. PubMed PMID: 22977183; PubMed Central PMCID: PMCPMC3592446.

34. Vlahovic I, Gluncic M, Rosandic M, Ugarkovic E, Paar V. Regular Higher Order Repeat Structures in Beetle Tribolium castaneum Genome. Genome Biol Evol. 2017;9(10):2668–80. doi: 10.1093/gbe/evw174. PubMed PMID: 27492235; PubMed Central PMCID: PMCPMC5737470.

35. Rosandic M, Paar V, Basar I, Gluncic M, Pavin N, Pilas I. CENP-B box and pJalpha sequence distribution in human alpha satellite higher-order repeats (HOR). Chromosome Res. 2006;14(7):735–53. doi: 10.1007/s10577-006-1078-x. PubMed PMID: 17115329.

36. Levenshtein VI. Binary codes capable of correcting deletions, insertions, and reversals. Doklady Akademii Nauk SSSR. 1965;163 (4):845–8.

